# Exploring the Impact of Probiotic Route of Administration on Its Protective Effects against Pathogenic Infection in *Galleria mellonella*

**DOI:** 10.1101/2024.01.20.576486

**Authors:** Lakshmi Naga Kavya Menta, Soffi Kei Kei Law, Hock Siew Tan

## Abstract

In probiotic research, identifying promising candidates for further development remains challenging. Traditional mammalian models are invaluable for assessing efficacy, but limitations such as high cost, ethical considerations, and lengthy reproductive cycles can slow the preliminary screening process. The *Galleria mellonella* larvae have emerged as a powerful *in vivo*-like screening model. Most studies on *G. mellonella* often use intra-hemocelic injection, bypassing the natural gut entry point for probiotics, which is analogous to introducing probiotics into the mammalian blood. Despite their advantages, discrepancies exist between *G. mellonella* and mammalian models, particularly regarding the route of probiotic administration. This study bridges this gap by investigating the differential effects of oral and intra-hemocelic administration of a probiotic, *Escherichia coli* Nissle 1917 (EcN1917), on its protective efficacy against a gastrointestinal pathogen, *Salmonella* Typhimurium ATCC 14028 (ST14028). The larvae were challenged with varying doses of EcN1917 and ST14028 through both routes. Survivability, health index, and bacterial burden in the gut and hemolymph were assessed. This study demonstrated that oral EcN1917 pre-treatment significantly increased survivability against ST14028 infection compared to the control group and alleviated the pathogen gut burden. Notably, injection pre-treatment decreased survivability. This discrepancy is attributed to the dual nature of probiotics, exhibiting beneficial effects in the gut but acting as pathogens in non-native locations like the hemolymph. This study concludes that the route of probiotic administration in *G. mellonella* significantly impacts the protective effects of probiotics.

## 1. Introduction

Traditionally, murine models are often used for the *in-vivo* study of probiotic intervention in managing microbial infections. *Galleria mellonella* larvae have recently served as ideal surrogate models for high-throughput studies [1] to overcome some associated ethical, economic and logistical restrictions, often slowing research progress. Although not as genetically well established as *Drosophila melanogaster* or *Caenorhabditis elegans*, the ease of manipulation, survivability at incubation temperatures of 25 to 37 °C and similarity of larval innate immune systems to mammalian models increase the attractiveness of this model system for studying protective effects of probiotics and the virulence of pathogens in a human physiological environment [2]. With the rapid rise in antimicrobial resistance in many pathogens [3][4], probiotics have been proposed as potential alternative antibiotics.

Probiotics are live, non-pathogenic microorganisms that benefit the host when consumed in appropriate oral doses and reside in the host’s gut after consumption [5]. However, most probiotics screening studies using *G. mellonella* have been conducted via intra-hemocelic injection, whereby the probiotics are introduced into the larval hemolymph, which is analogous to introducing probiotics into the mammalian blood [5][6][7]. This approach is misleading and might give an inaccurate interpretation of the protective effects of the tested probiotics. Therefore, our study aims to investigate and compare the differences in the degree of protection conferred by a probiotic against a gastrointestinal pathogen via different routes of administration (force-feeding vs injection).

In this study, *Escherichia coli* Nissle 1917 (EcN1917), a commercialised and well-characterised therapeutic with over a century-long history of human use [8], was used as the model probiotic while *Salmonella* Typhimurium, a widespread gastrointestinal pathogen with increasing outbreaks of drug-resistant infections [9] was used to establish a gut infection of the larval model. This study demonstrated that the route of probiotic administration significantly impacts its protective effects and should be considered when screening for probiotics in *G. mellonella*.

## 2. Methods and Materials

### 2.1 Bacterial strains, culture preparations and growth curves

Probiotic EcN1917 and pathogenic ST14028 were cultured and routinely maintained on nutrient agar. Growth kinetics for each bacterial strain were determined to prepare active bacterial suspensions with the highest concentration of viable cells for larval inoculation. Concisely, the bacterial overnight cultures were subcultured in Luria Bertani (LB) broth and incubated at 37°C, shaking at 200 rpm. The optical density (OD 600 nm) was determined every 30 minutes for 6 hours until consistent consecutive measurements were obtained. The spectrophotometric measurements from three replicates were used to plot growth curves and determine the respective lag, exponential and stationary phases.

### 2.2 G. mellonella challenge assays

Batches of *G. mellonella* larvae were sourced from Carolina Biological, USA and maintained as per established protocol [10]. Larvae without melanisation were incubated overnight and acclimatised to 37 °C before use in the challenge assays. Bacteria were grown to the mid-log phase, harvested by centrifugation (6700g, 7 minutes) and washed to remove traces of media or compounds produced *in vitro*. They were serially diluted 10-fold using 1X PBS [10]. Larvae were sterilised using 70% ethanol and injected or force-fed with 10μL of bacterial inocula using a Hamilton syringe. For force-feeding, blunt-end needles were used to prevent internal punctures. Two additional control groups were included: a PBS-only group accounting for physical trauma caused by the inoculation process and a no-inoculation group accounting for natural deaths. The larvae were incubated at 37° C; melanisation and survival were monitored and recorded at 0-, 3-, 6-, 24-, 48-, 72-, 96-, 120- and 144 hours post-inoculation. The amount of bacteria (colony-forming units: CFU/mL) administered was determined by plating the prepared inocula on LB agar. The assays were performed in duplicates for each bacterial strain, with 5 larvae per treatment group, via force-feeding and injection.

### 2.3 G. mellonella prophylactic protection assays

Bacterial inocula (probiotic EcN1917 and pathogen ST14028) were prepared according to section 2.3. Larvae were inoculated according to treatment groups (as specified in Table 1) and incubated at 37°C for 2 hours before the second inoculation. Larval survival, activity, and melanisation were observed and scored at various time points post-second inoculation. Protection assays were performed in triplicates for both injection and force-feeding techniques.

**Table 1.**
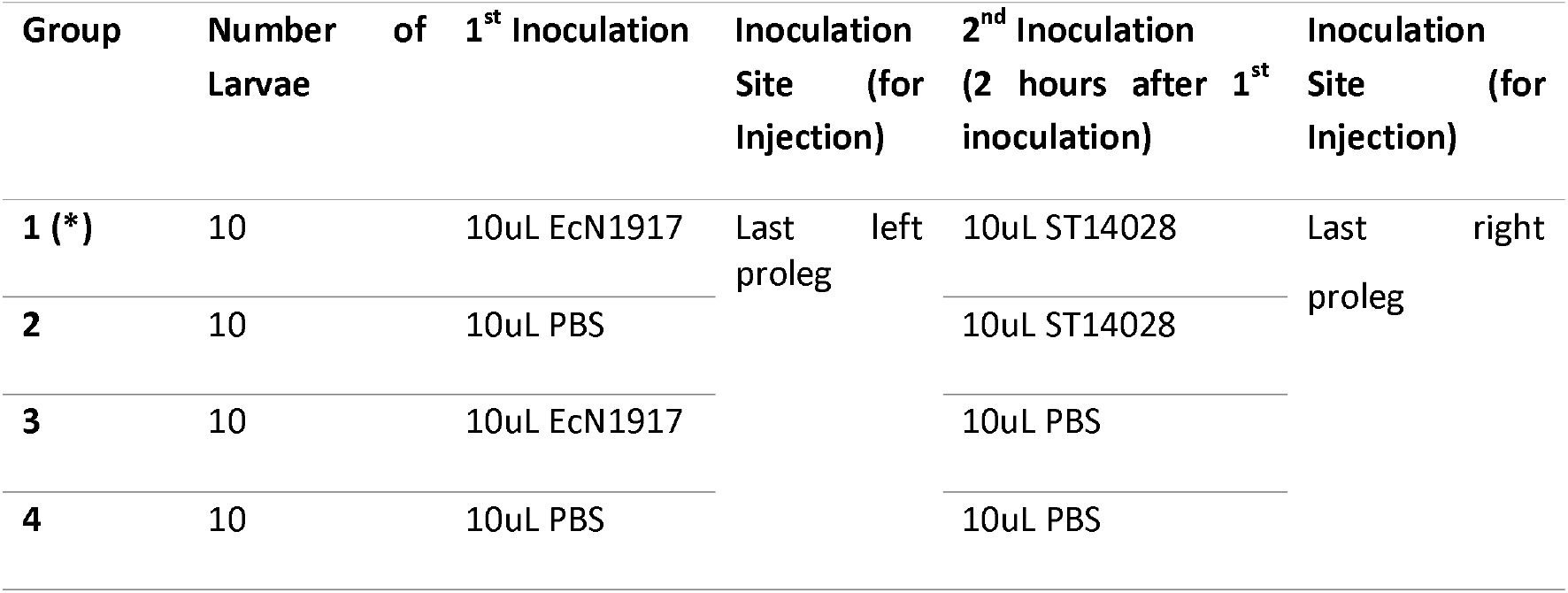
Experimental groups were used for the protection assays. Group 1* represents the treatment group (pre-treated with probiotics before infection with the pathogen). Groups 2 to 4 were the infection-only, probiotic-only and trauma controls, respectively.

### 2.4 Bacterial transformation and G. mellonella bacterial burden assays

EcN1917 and ST14028 were transformed using a heat-shock approach with selection plasmids. Chemically competent ST14028 and EcN1917 cells were transformed with pULTRA_RFP_KAN and pULTRA_CmR_KAN, respectively. Briefly, the larvae were treated with transformed EcN1917 and ST14028 following the protection assay protocol. At 0-, 4-, and 24 hours post-inoculation, the hemolymph and gut were extracted and dissected from 2 randomly selected larvae (24 hours was chosen as the final time point to ensure possible extraction). To release intracellular bacteria, 1μL of 5mg/mL Digitonin was added to the extractions, followed by centrifugation (2300g, 4 °C, 7 minutes) [10]. The bacterial supernatants were serially diluted and plated on different selective agars. 50 μg/mL kanamycin (KAN) and RFP expression were used for ST14028 selection. Cell extracts were plated on agar containing 50 μg/mL KAN and 15 μg/mL chloramphenicol (CHL) to select for EcN1917. Plates were incubated at 37° C, and the number of CFU was determined and normalised to the weight of the larval extractions. Assays were carried out in triplicates.

### 2.5 Statistical Analysis

Results from replicates of experiments were pooled and statistically analysed using GraphPad Prism 8 (GraphPad Software, La Jolla, USA). Kaplan-Meier survival curves were plotted for the challenge and protection assays, and the significance differences were predicted using the Mantel-Cox log-rank tests (significance at p ≤ 0.01). Multiple t-tests (significance at p ≤ 0.05) were conducted to compare the differences between health index scores and bacterial burden presented in clustered bar graphs. Statistical significance between two groups is visualised as asterisks in figures (⍰p ≤ 0.05; ⍰⍰p ≤ 0.01; ⍰⍰⍰p ≤ 0.001; ⍰⍰⍰⍰p ≤ 0.0001).

## 3. Results and Discussion

### 3.1 *Route of probiotic and pathogen administration alter* G. mellonella *survivability*

Both oral force-feeding (Figure 1a) and intra-hemocelic injection (Figure 1b) were tested in the challenge assays to identify the difference in potencies of the bacterial strains. Dose-response curves were plotted for each bacterial strain and inoculation route using data from the 72-hour time point by which maximal deaths were observed. While injections caused a concentration-dependent decline in survivability at lethal dose LD50 of 10^6^ CFU/mL (Figure 1c), larval survivability remained at 100% after force-feeding with all concentrations (10^5^ to 10^8^ CFU/mL) of EcN1917 (Figure 1d). While probiotics are safe in the gut, positively modulating host microbiota and immune responses following oral administration [7], they can cause disease in systemic circulation. Probiotic bacteraemia, though rare, has been documented in immunocompromised individuals [11][12]. Similarly, EcN1917, despite being considered safe, was found to harbour the *pks* pathogenicity island, encoding non-ribosomal and polyketide synthases involved in colibactin production [13]. This genotoxic factor triggers DNA damage during systemic infection. Hence, introducing EcN1917 into the larval hemolymph via injection may cause both bacteraemia and septicaemia, contributing to larval mortality at higher concentrations [13, 14]. This observation underscores the dynamic nature of non-pathogenic EcN1917 responses depending on the inoculation site.

**Figure 1.**
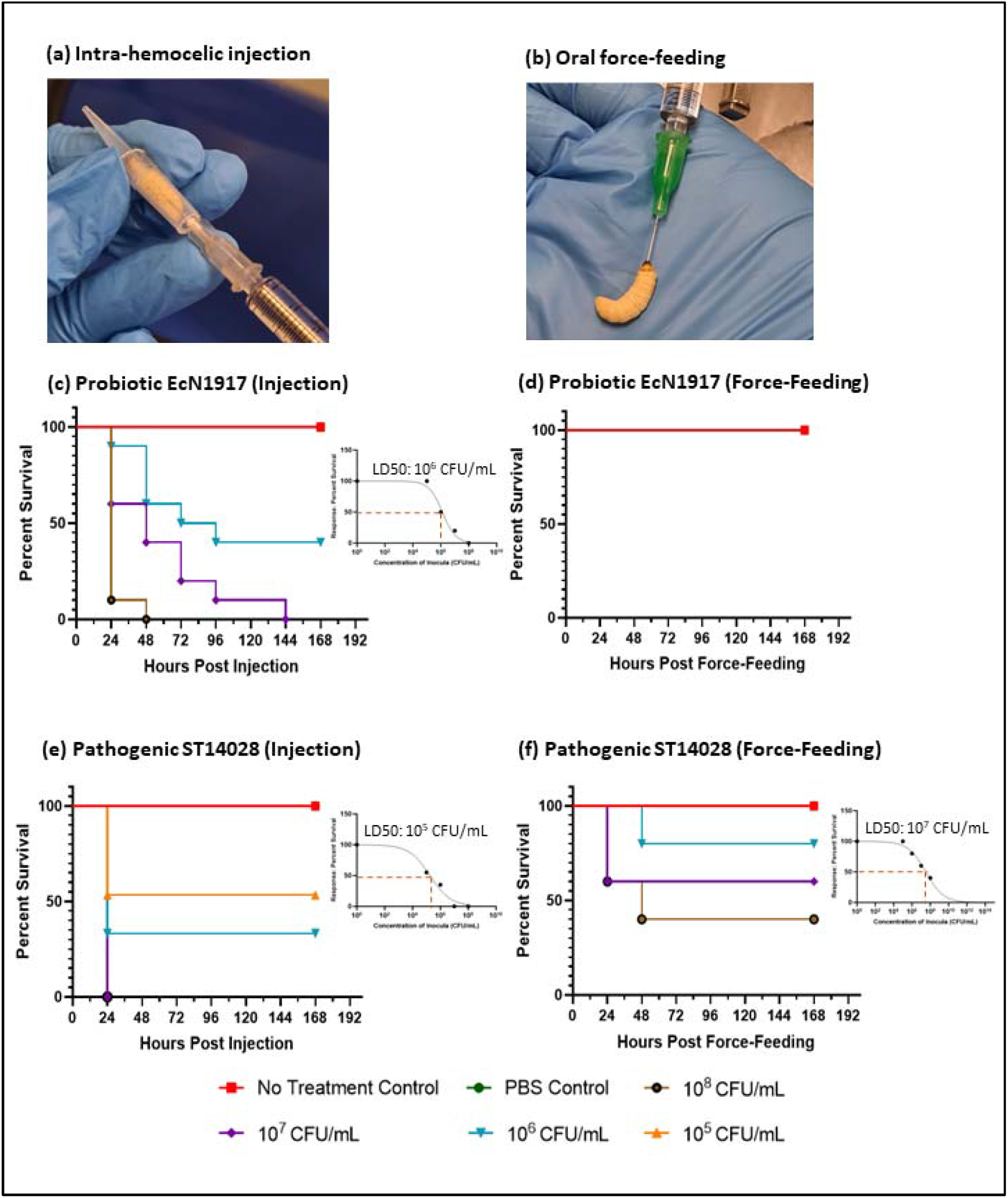
(a) Larval inoculation via intra-hemocelic injection. (b) Larval inoculation via oral force-feeding. Kaplan Meier Survival curves showing the effects of inoculating various concentrations of the EcN1917 via (c) Injection (d) Force-feeding (survival remained at 100% in treatment groups) and ST14028 via (e) Injection (f) Force-feeding. All differences in survivability between treatment groups were significant (p-value < 0.01), Mantel-Cox log-rank test). (n=5 per treatment group, 2 Biological replicates). Dose-response curves, with the lethal dose to 50% of the larval population, have been presented.

In contrast, inoculation with ST14028, an enteric pathogen, decreased larvae survivability through both routes of inoculation (Figure 1e and 1f). Notably, the larvae were more susceptible to ST14028 when injected intra-hemocelically (LD50 of 10^5^ CFU/mL) than force-fed into the gut (LD50 of 10^7^ CFU/mL). The LD50 values indicate that a lower dosage of the pathogen ST14028 was required to cause larval death following injection. The presence of pathogens in the larval hemolymph triggers the immune cells to produce reactive oxygen species (ROS) to eradicate them. This immune response mechanism mirrored that in mice models with systemic *S*. Typhimurium infection [15]. When infected with high doses of pathogens, elevated hemolymph ROS levels are detrimental to the larvae [16], which aligns with our observation that higher ST14028 inocula concentrations increase larvae deaths. The eicosanoids in *G. mellonella* possibly lose functionality at high ROS levels, disrupting anti-oxidative enzymes, leading to ROS accumulation and increased oxidative stress, inducing lipid peroxidation and immune cell cytoskeleton disruption, damaging essential proteins that further deteriorate larval health [17]. Impacts of *S*. Typhimurium in the larvae parallel observations in humans, where infections affecting the systemic circulation elevate mortality risk and severity of symptoms compared to those limited to the gastrointestinal tract [18]. These findings underscore the importance of selecting an appropriate larval inoculation route to closely replicate mammalian models’ internal physiological conditions and increase the relevance of generated results to practical applications.

### 3.2 Oral administration of Ecn1917 conferred protection against ST14028 infection

The larvae were pre-treated with the probiotic and then infected with the pathogen 2 hours later to study the prophylactic effects of EcN1917 against ST14028. Their survivability and health were monitored using a previously established *G. mellonella* Health Index Scoring [19] (Figure 2a). The maximum score of 7 indicated healthy larvae with no physical signs of melanisation or decline in activity. Based on the results from the challenge assays, 10^5^ CFU/mL (injection) and 10^6^ CFU/mL (force-feeding) of both bacteria were selected for the protection assays. These doses of probiotic did not harm the larvae yet maintained the pathogen’s infectivity to cause larval death.

**Figure 2.**
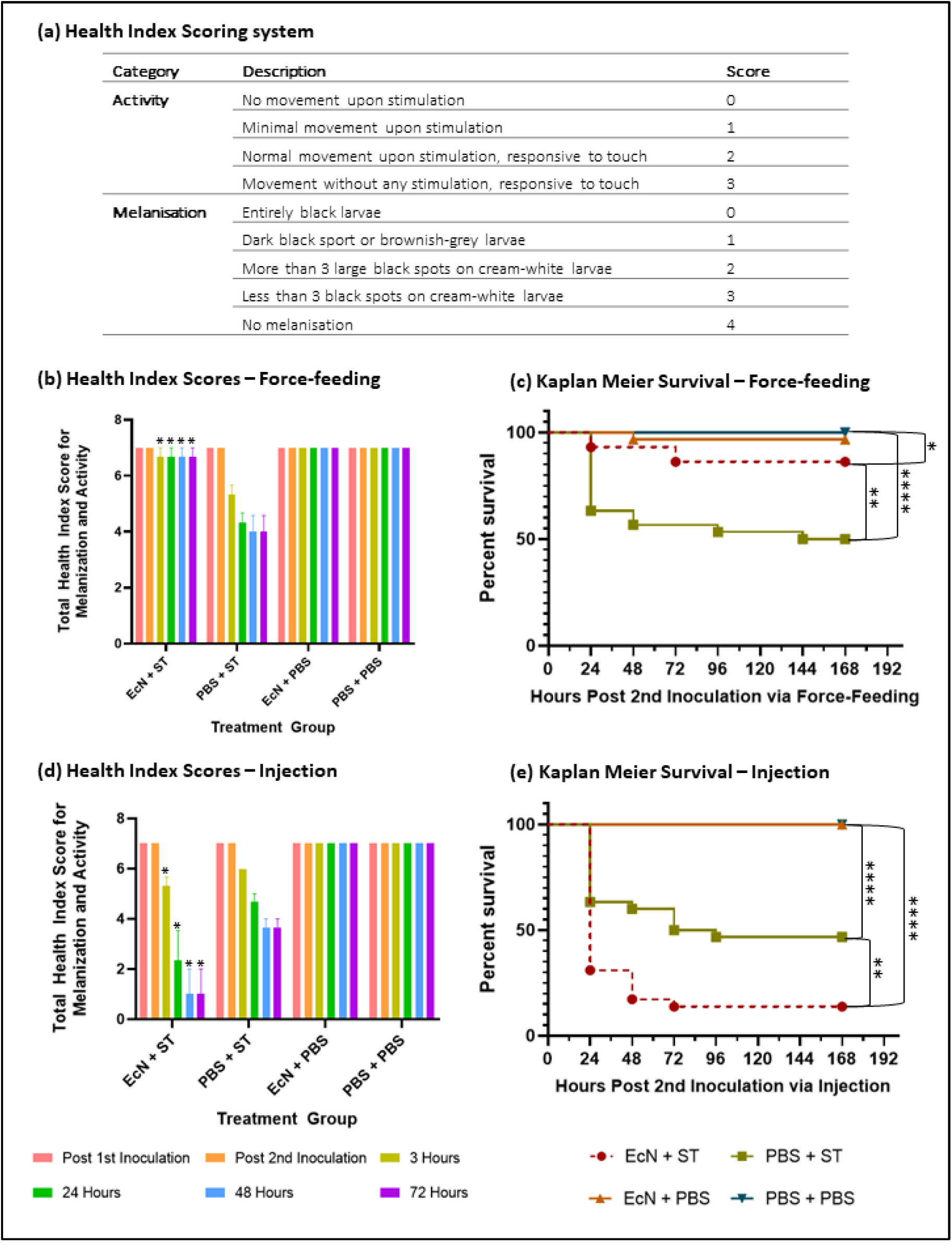
(a) Health Index Scoring system for monitoring larval health. (b) Health Index Scores for first 3 days via force-feeding (c) Kaplan Meier Survival curve for protection assay via force-feeding (d) Health Index Scores for first 3 days via injection (c) Kaplan Meier Survival curve for protection assay via injection. All differences in survivability between treatment groups were significant (p < 0.01, Mantel-Cox log-rank test) (n=10 for each treatment group, 3 Biological replicates). Health scores of the treatment group (EcN + ST), which were significantly different (p < 0.05) from the infection-only group (PBS + ST), have been indicated (*).

The differences in larval health index scores between the force-fed EcN + ST and PBS + ST were significant at 3, 24, 48 and 72 hours (p < 0.05), with higher mean health scores for the pre-treated group (Figure 2b). Concordantly, the larvae survivability dropped to 50% by day 7 in the ST14028 infection-only group, while the larvae group that received prophylactic pre-treatment with EcN1917 showed elevated survivability to 85% (p-value 0.0027) (Figure 2c).

Conversely, probiotic pre-treatment administered through injection lowered larval survivability to 15%, compared to injection with pathogen alone, which saw 45% survival (p-value 0.0024). The EcN + ST group had lower total health scores than the PBS + ST group at 3, 24, 48 and 72 hours (Figures 2d and 2e). The health scores of the treatment group were also significantly lower than the controls (EcN + PBS and PBS + PBS). These findings corroborated with the challenge assays, where the probiotic could induce larval death when injected alone (Figure 1c), and the larvae faced an elevated risk of mortality when the pathogen was injected rather than force-fed (Figure 1e and 1f). The synergy in larval mortality with probiotic pre-treatment and pathogen infection via injection suggests probiotics cannot exert protective effects if administered into hemolymph instead of the gut of *G. mellonella*.

Rossoni et al. revealed that injection of some probiotic *Lactobacillus paracasei* strains, although capable of prolonging the life of larvae challenged with *Candida albicans*, was incapable of increasing the percentage survivability by the end of the experimental period [5]. In other words, the rate at which the fungal infection caused death was slowed down. Similarly, Jorjão et al. demonstrated that pre-treatment of larvae with *Lactobacillus rhamnosus* via injection decreased the survivability of larvae infected with *Staphylococcus aureus* or *E. coli* compared to those that were not pre-treated [7]. The findings from these two authors are consistent with this study, suggesting that inoculation of probiotics into the larval hemolymph should be avoided as it may not reflect the real protective effects of probiotics.

### 3.3 Prophylactic pre-treatment of larvae with EcN1917 decreases the ST14028 burden

Bacterial burden assays were conducted using transformed EcN1917 (EcN1917* carrying a chloramphenicol selectable marker in a plasmid) and ST14028 (ST14028* carrying a Kanamycin selectable marker and red fluorescence protein in a plasmid) to ascertain the correlation between the observed protective effects of EcN1917 with the quantities of ST14028 and EcN1917 within the larvae (Figure 3a). The virulence of EcN1917* and ST14028* was similar to the corresponding untransformed strains (Figure 3b), indicating that the introduction of plasmids carrying the antibiotics’ selectable marker had no impact on bacterial virulence and can be used for subsequent bacterial burden experiments.

**Figure 3.**
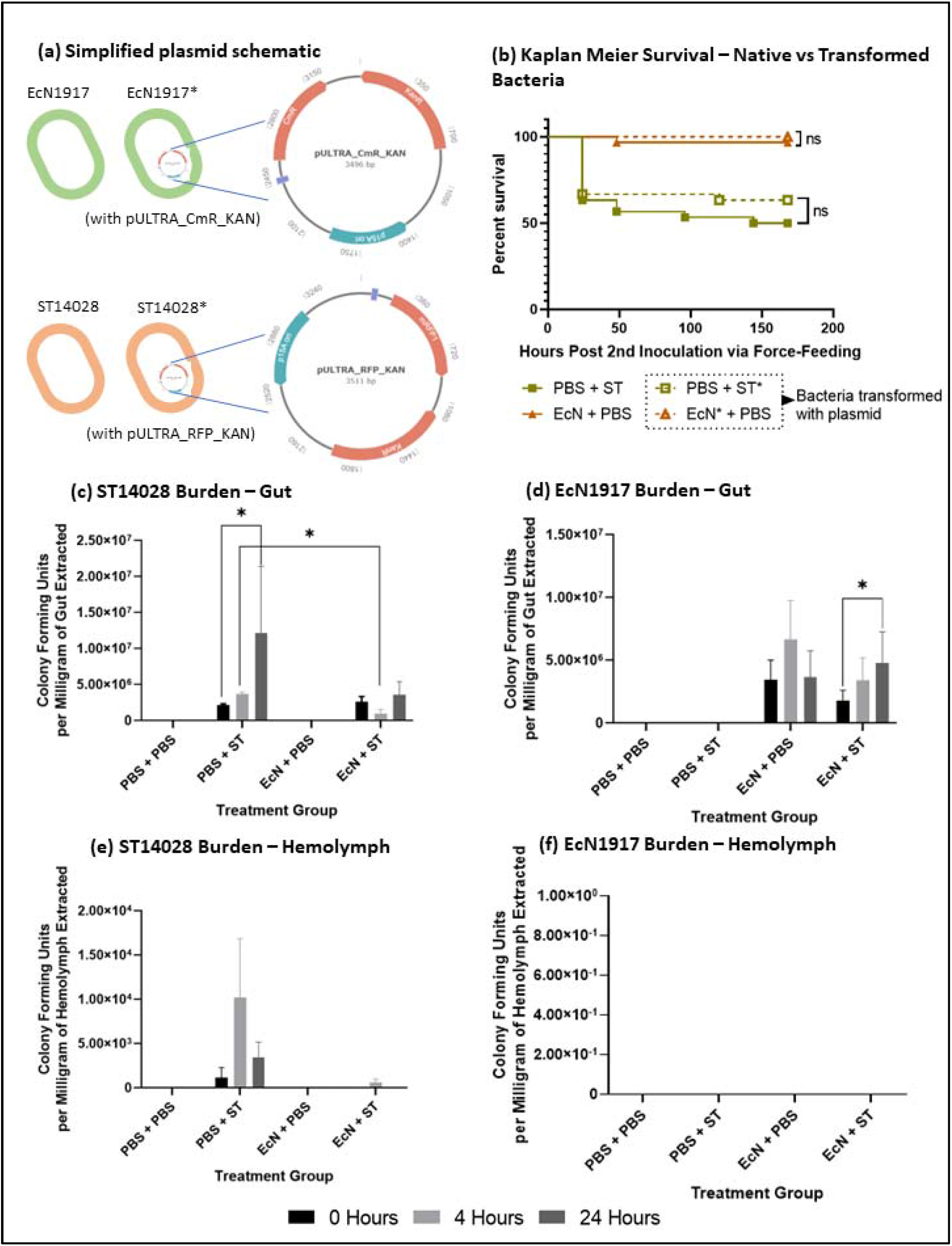
(a) Simplified schematic of plasmids with selection markers used for enumeration of EcN1917 and ST14028 (b) Kaplan Meier Survival curve comparing native and transformed(*) EcN1917 and ST14028 (c) ST14028 burden in the larval guts (d) EcN1917 burden in the larval guts (e) ST14028 burden in larval hemolymph (f) EcN1917 burden in the larval hemolymph.

The bacterial burden of ST14028 in the gut (Figure 3c) of the infection-only group (PBS + ST) was higher than the probiotic pre-treated group (EcN + ST) at all time points, with a notable difference at 4 hours (p-value 0.012). The pathogen numbers also increased drastically between 0 and 24 hours without probiotic pre-treatment (p-value 0.034). Conversely, the pathogen burden in the probiotic-treated group remained almost consistent throughout the 24-hour experimental period. Moreover, no significant differences in probiotic abundance within the gut were observed between the treatment groups (EcN + ST and EcN + PBS) (Figure 3d). However, the levels of EcN1917 in the gut of the larvae pre-treated with EcN1917 before the ST14028 challenge were significantly increased (p-value 0.030).

The reduced pathogen abundance and the concurrent increase in probiotic levels observed with probiotic pre-treatment can be attributed to EcN1917’s capacity to limit the availability of micronutrients, such as iron, which are crucial for *Salmonella* proliferation and pathogenicity [20] [21]. The availability of free iron in healthy larvae is naturally low due to iron homeostasis by ferritin and transferrin [22]. During infections, hemocytes produce lipocalin, a protein that reduces free iron levels, while sequestering iron from the pathogen’s siderophores. The introduction of probiotic EcN1917 further aggravates the stringency of iron availability *in-vivo* by intercepting iron-bound *Salmonella* siderophores and concurrently producing an array of its siderophores competing with the pathogen for free iron uptake [21][23].

Besides, an *in-vitro* study reported that EcN1917 could produce microcin MccH47 exhibiting antibacterial effects against *S*. Typhimurium, which is enhanced in iron-limited niches [24]. Azpiroz and Lavina [25] stated that in the presence of pathogens, EcN1917 produces pre-microcin MccH47, which promotes an increase in enterobactin siderophore expression. This event triggers the post-translational siderophore moiety attachment to the pre-microcin. MccH47 then mimics iron-siderophore complexes and is taken up by the pathogen, leading to unregulated proton entry and causing cell lysis [26].

The hemolymph pathogen burden in untreated larvae was higher than treated at all time points. Minimal levels of ST14028, approximately 10^2^ CFU/mg, were detected in the hemolymph of untreated larvae at 4 hours post-infection (Figure 3e), but this did not lead to larval melanisation or significant deaths by 24 hours. There was no significant difference in the health score for the probiotic pre-treated (EcN + ST) group compared to the controls (EcN + PBS and PBS + PBS) 24 hours post-2nd inoculation (Figure 2c).

In contrast, more than 10^4^ CFU/mg of ST14028 was present in the larval hemolymph for the group infected with pathogen ST14028 only (PBS + ST) at 4 hours post-infection (Figure 3e). This group’s survivability and health scores dropped to 60% and 4 by 24 hours (Figures 2c and 2d). This suggests that a minimum number of pathogens are required to activate a strong antimicrobial immune response with visible melanisation. Bismuth et al. reported a minimum of 10^4^ CFU/larvae of *S*. Typhimurium had to be present within the larval hemolymph to cause death [27].

## Conclusion

This study revealed dose- and route-dependent effects of probiotic EcN1917 and pathogenic ST14028 in *G. mellonella* larvae. Injection with probiotics led to significant larval mortality and health decline at concentrations above 105 CFU/mL, while force-feeding showed no adverse effects. The protection assays demonstrated enhanced survival with oral pre-treatment, contrasting the decline in survivability through injection. This work is the first probiotic protection study in *G. mellonella* larvae that has reported the direct comparison between outcomes from oral and intra-hemocelic inoculation. Future research could explore the possible mechanisms by which the probiotic enhances the intestinal gut barrier and resists pathogen colonisation, emphasising the importance of route and dose of probiotic *G. mellonella* larval administration.

## Contributors

Kavya Menta: investigation, formal analysis, writing - original draft, review & editing, visualisation

Soffi Law: writing: review & editing

Hock Siew Tan: supervision, conceptualisation, writing - review & editing

## Acknowledgements

This study was supported by the School of Science, Monash University Malaysia.

